# Brain-wide screen of prelimbic cortex inputs reveals a functional shift during early fear memory consolidation

**DOI:** 10.1101/2022.03.16.484644

**Authors:** Lucie Dixsaut, Johannes Gräff

**Affiliations:** Laboratory of Neuroepigenetics, Brain Mind Institute, Ecole Polytechnique Fédérale de Lausanne (EPFL), Switzerland

## Abstract

Memory formation and storage rely on multiple interconnected brain areas, the contribution of which varies during memory consolidation. The medial prefrontal cortex, in particular the prelimbic cortex (PL), was traditionally found to be involved in remote memory storage, but recent evidence points towards its implication in early consolidation as well. Nevertheless, the inputs to the PL governing these dynamics remain unknown. Here, we first performed a brain-wide, rabies-based retrograde tracing screen of PL engram cells activated during contextual fear memory formation to identify relevant PL input regions. Next, we assessed the specific activity pattern of these inputs across different phases of memory consolidation, from fear memory encoding to recent and remote memory recall. Using projection-specific chemogenetic inhibition, we then tested their functional role in memory consolidation, which revealed a hitherto unknown contribution of claustrum to PL inputs at encoding, and of insular cortex to PL inputs at recent memory recall. Both of these inputs further impacted how PL engram cells were reactivated at memory recall, testifying to their relevance for establishing a memory trace in the PL. Collectively, this data identify a spatiotemporal shift in PL inputs important for early memory consolidation, and thereby help to refine the working model of memory formation.

## Introduction

The brain’s ability to form enduring memories is essential for an individual’s survival. Memories first need to be encoded and subsequently stored in the brain, a process that is termed memory consolidation. Memory consolidation occurs both at the scale of individual cells (cellular consolidation), which happens in the order of seconds to hours, and at the scale of brain networks (systems consolidation), which takes place in the days to weeks after learning (Dudai, 2004, 2012). Systems consolidation across several brain areas is thought to be essential for the establishment of enduring memories (Nadel & Hardt, 2011).

Traditionally, the hippocampus (HPC) was demonstrated to be necessary during the early stages of memory formation and the retrieval of recent memories (which in the mouse are typically studied one day after encoding), while the medial prefrontal cortex (mPFC) was found to be rather responsible for the consolidation and retrieval of remote memories (which are studied at least 14 days after encoding) (Albo & Gräff, 2018; Frankland & Bontempi, 2005). Indeed, multiple studies using immediate early gene mapping (IEGs, that are expressed specifically upon neuronal activation), whole brain region inhibition or cell type specific optogenetic manipulations (Aceti et al., 2015; Frankland et al., 2004, 2006; Goshen et al., 2011; Makino et al., 2019; Silva et al., 2018; Wheeler et al., 2013) showed that the mPFC was predominantly important at later times as opposed to the HPC. However, recent evidence has challenged this view by highlighting a role for the mPFC also during fear memory encoding (Bero et al., 2014; Cho et al., 2017; Cummings & Clem, 2020; Tang et al., 2005; Zelikowsky et al., 2013) as well as for fear memory recall at recent times (Do-Monte et al., 2015; Rajasethupathy et al., 2015). The rich connectivity of the mPFC indeed places it as a potential hub region for memory consolidation, as it receives not only inputs from other cortical areas, including sensory ones, but also from various subcortical areas such as the hippocampal formation, amydgala and thalamus (Dixsaut & Gräff, 2021; Le Merre et al., 2021), which are all implicated in memory formation (Cho et al., 2017; Nonaka et al., 2014; Ramirez et al., 2013; Reijmers et al., 2007; Taylor et al., 2021).

At the cellular level, mounting evidence suggests that memories are encoded and stored in *engram* cells, which, by definition (Tonegawa et al., 2015), are cells that are activated during the initial learning, undergo molecular and/or cellular modifications following learning, and the reactivation of which correlates with and can trigger memory recall. Engram cells have been discovered not only in the HPC (Josselyn et al., 2015; Liu et al., 2012; Ramirez et al., 2013), but more recently also in the mPFC (DeNardo et al., 2019; Kitamura et al., 2017; Matos et al., 2019). Interestingly, engram cells in the mPFC were reported to have the particular feature of staying silent until the memory is fully consolidated, although they are formed during the original learning phase (DeNardo et al., 2019; Kitamura et al., 2017; Matos et al., 2019). This implies that mPFC engram cells are first active during encoding, stay silent during a recent recall, and are reactivated at remote recall, although they are functionally able to trigger memory retrieval at any time.

Based on these grounds, we hypothesized that the functional contribution of mPFC inputs may change over the course of memory consolidation to govern how the mPFC engram is formed and subsequently reactivated. For this reason, we first sought out to establish a comprehensive functional map of mPFC inputs across time during fear memory consolidation, and second to analyze the downstream effect of these inputs on memory retention and mPFC engram reactivation.

## Results

### The prelimbic cortex is specifically active during the encoding of a fear memory

The mPFC is composed of the three following major areas: The anterior cingulate (ACC), the prelimbic (PL) and the infralimbic (IL) cortices (Carlén, 2017; Le Merre et al., 2021). In order to evaluate the relative activity of these subregions during the different phases of fear memory consolidation, we used a contextual fear conditioning (CFC) paradigm in combination with cFos immunohistochemistry (IHC), an IEG marker of neuronal activity. We measured the freezing percentage of wild-type (WT) mice at CFC encoding before the footshocks occurred, as well as at recent recall 1 day post encoding and at remote recall 14 days post-encoding. Each group was controlled for by a no shock group. We observed that both at recent and remote recall, mice display a significantly higher freezing percentage than the no shock control groups (Figure 1B), indicating successful memory formation.

**Figure 1.**
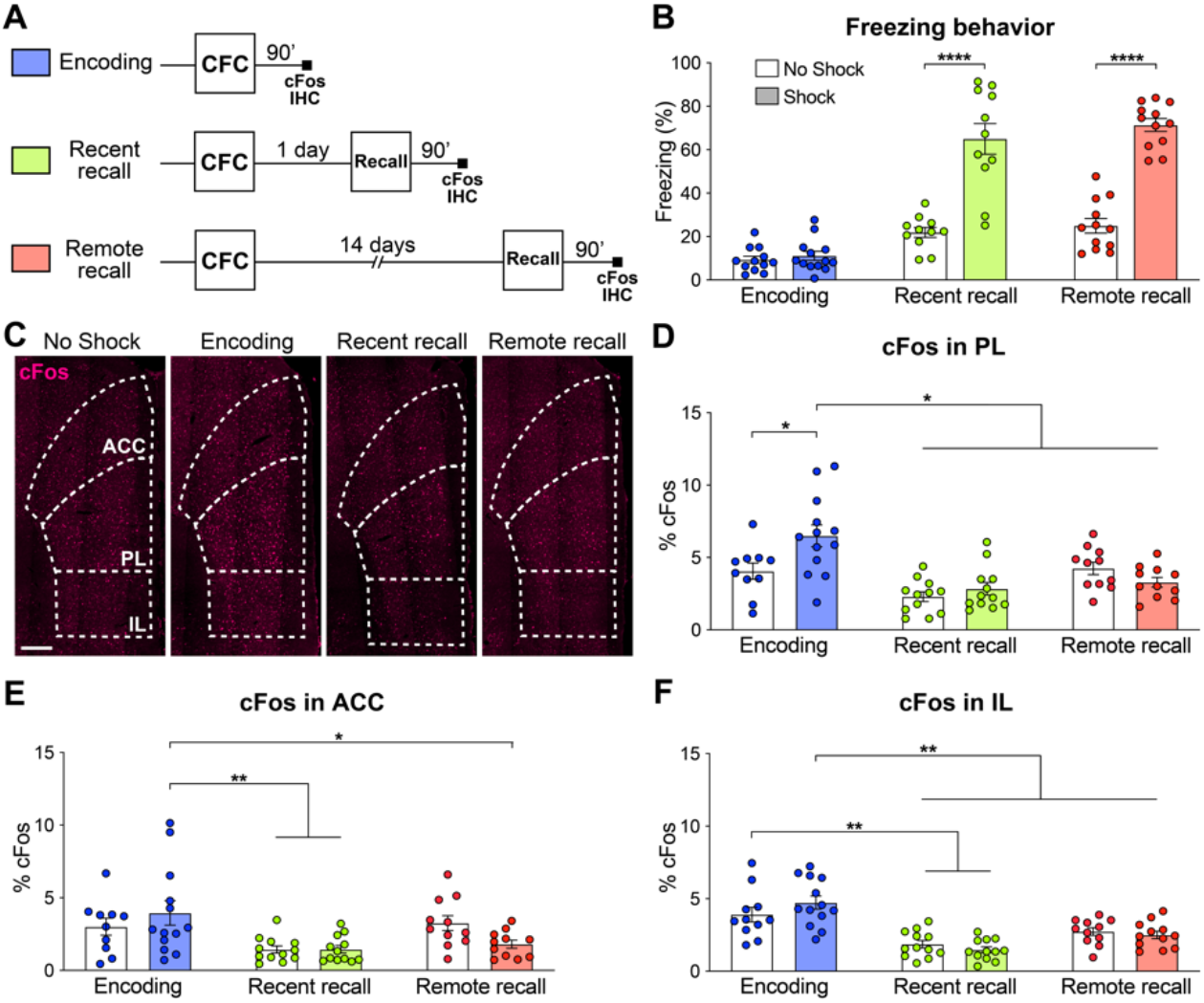
The prelimbic cortex (PL) is activated by the encoding of a contextual fear memory. **(A)** Experimental design. For encoding, mice were perfused 90 minutes after contextual fear conditioning (CFC). For recent and remote recalls, mice were perfused 90 minutes after a 1-day and a 14-day recall, respectively. **(B)** Percentage freezing measured during the 3 minutes of habituation before the shocks (Encoding, in blue), at recent (in green) or remote (in red) memory recalls, for the animals undergoing CFC (shock, filled) and the control groups that were exposed to the CFC chamber without the shock (no Shock, clear). Two-tailed unpaired t-tests, ****: p < 0.0001. **(C)** Representative images of cFos immunostainings in the mPFC. Scale: 250μm. **(D-F)** Percentage of cFos over DAPI in **(D)** PL (one-way-ANOVA, F (5, 63) = 9.172, p < 0.0001), **(E)** ACC (one way ANOVA, F (5, 63) = 4.394, p = 0.0017) and **(F)** IL (one way ANOVA, F (5, 65) = 13.34, p < 0.0001). **(D-F)** Stars represent least significant p-values of Tukey’s multiple comparisons tests: *: 0.01<p<0.05; **: 0.001<p<0.01. n = 11-13 animals per group.

We then quantified cFos expression in the three subregions of the mPFC (Figure 1C) 90min after the corresponding behavioral session. We found that while all regions were more active at encoding than during the memory recalls, only in the PL did we observe a significant increase in cFos compared to the no shock control groups (Figure 1C-F). These results indicate that the mPFC as a whole, and the PL in particular, are activated by the encoding of a contextual fear memory. In turn, this finding suggests an important role of the PL already during this early phase of memory consolidation, which is coherent with the formation of engram cells in the PL at the time of encoding (DeNardo et al., 2019; Kitamura et al., 2017; Matos et al., 2019).

### Brain-wide screen of PL engram inputs

Next, we aimed to identify the PL inputs that could be responsible for this peaked PL activity at encoding and for the establishment of its engram during memory consolidation (DeNardo et al., 2019; Kitamura et al., 2017; Matos et al., 2019). To this end, we employed an activity-dependent monosynaptic retrograde tracing technique (Figure 2A,B). Specifically, we used the TRAP2 mouse line (DeNardo et al., 2019), in which the cFos promoter drives the expression of the tamoxifen-dependent Cre^ERT2^ recombinase. These mice were first injected in the PL with helper AAVs expressing Cre-dependent nuclear GFP, the TVA receptor and optimized rabies glycoprotein oG. Thus, the expression of these proteins in PL engram cells could be triggered with tamoxifen injection at the time of encoding. 3 weeks post-encoding, we injected a modified rabies vector RVΔG-mCherry (Wickersham et al., 2007) that can only infect and replicate in TVA- and oG-expressing cells, respectively, allowing us to transsynaptically label all monosynaptic inputs of PL engram cells (Figure 2B). As expected, we found that tamoxifen injection increased the number of starter cells expressing both GFP and mCherry in PL compared to vehicle (Figure 2C-E), confirming the specificity of these tools to restrict tracing to PL engram cells.

**Figure 2.**
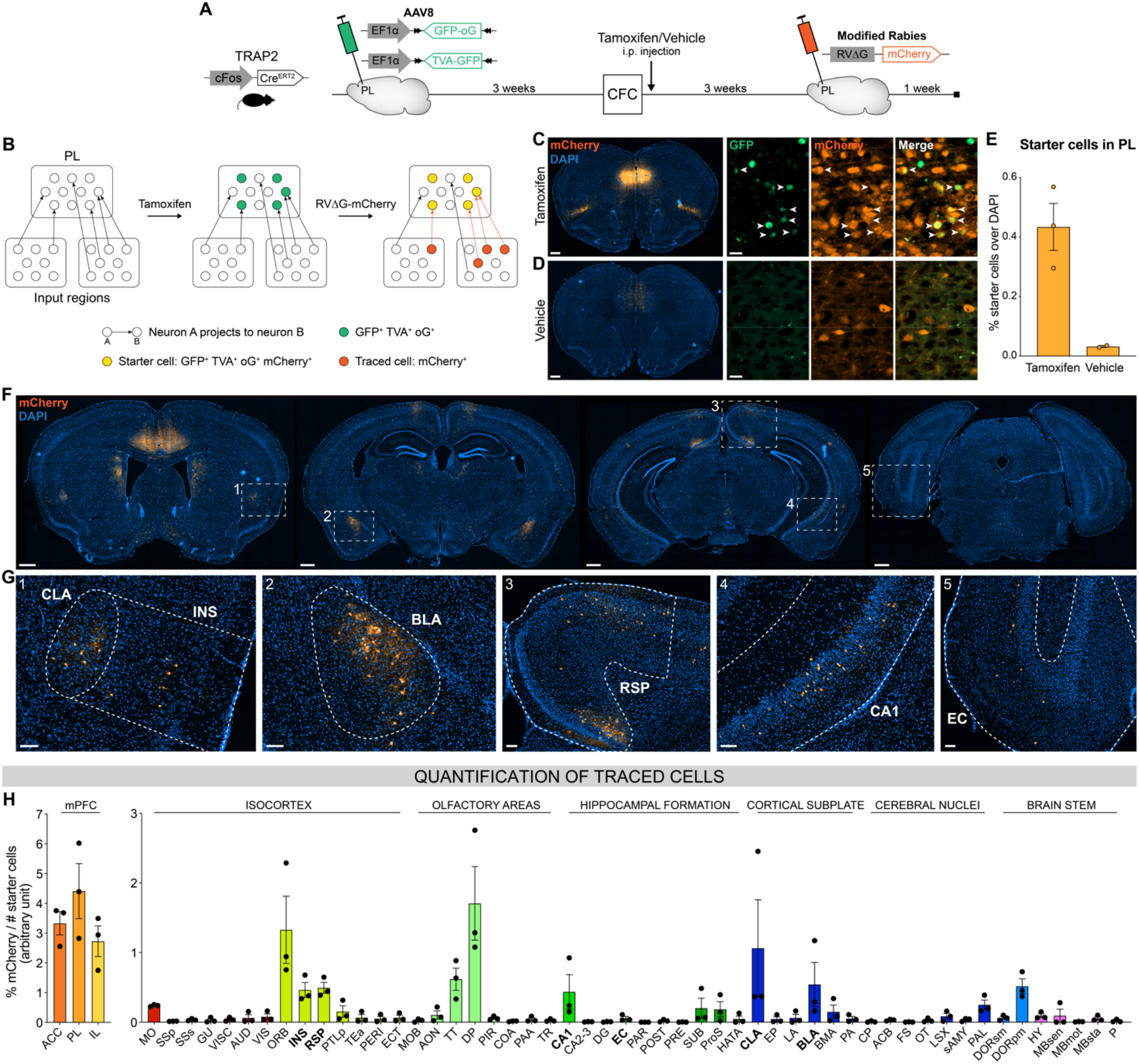
Brain-wide retrograde tracing identifies monosynaptic inputs of PL engram cells. **(A-B)** Experimental design and timeline. cFos-Cre^ERT2^ animals were first injected in the PL with helper AAVs expressing GFP, TVA receptor and oG (rabies optimized glycoprotein) in a Cre-dependent manner. Tamoxifen (or vehicle for control) was injected right after CFC to trigger recombination in cFos-Cre^+^ cells. 3 weeks later, a modified rabies virus (RVΔG-mCherry with EnvA coating) was injected in PL where it infected TVA-expressing cells, replicated in oG expressing-cells, and was retrogradely transsynaptically transported. A week later, brains were collected to quantify monosynaptic inputs of PL engram cells labelled with mCherry. **(C, D)** Representative images of the PL injection site (scale 400μm) and magnified view of starter cells (scale 20μm) with tamoxifen **(C)** or vehicle **(D)** injection. **(E)** Percentage of starter cells over DAPI in PL. **(F)** Representative images of traced cells throughout the brain (scale 500μm). **(G)** Magnified views of traced cells (scale 100μm) in CLA (inset 1), INS (1), BLA (2), RSP (3), vCA1 (4) and EC (5). **(H)** Brain-wide quantification of traced cells, normalized by the number of starter cells for each animal, in the mPFC subregions (left) and the rest of the brain (right). Tamoxifen: n = 3 animals; Vehicle: n = 2 animals.

We then quantified the percentage of traced cells (mCherry^+^ only) throughout the brain (Figure 2F-H). Although most traced cells were found in the mPFC itself and its neighboring areas (orbitofrontal cortex OFC and dorsal peduncular DP, Figure 2H), we observed traced inputs in several other brain regions, notably the claustrum (CLA, Figure 2G inset 1), insular cortex (INS, inset 1), basolateral amygdala (BLA, inset 2), retrosplenial cortex (RSP, inset 3), CA1 field of the HPC (mostly the ventral part, inset 4), taenia tecta (TT), thalamus polymodal association cortex related areas (DORpm), subiculum (SUB), and to a lesser extent the entorhinal cortex (EC, inset 5). Without tamoxifen injection, traced cells were negligible (Figure 2 S1), which further confirms the rabies tracing specificity.

With this approach, we identified relevant PL inputs that might be responsible for the development of the PL engram cell population, but we still lack information on whether, when and the extent to which these inputs are activated across memory consolidation.

### PL inputs are differentially activated across memory consolidation

Out of the regions projecting to PL engrams, we selected six brain areas with consistent input tracing for further investigation, because of their previously documented implication in various aspects of fear memory: The EC, for its role in memory formation (Roy et al., 2017) and its known projection to the mPFC necessary at encoding (Kitamura et al., 2017) and retrieval (Pilkiw et al., 2022); the RSP for its necessity for recent (Cowansage et al., 2014) and remote fear memory recall (Todd et al., 2016); the INS for its requirement during the consolidation and expression of contextual fear memories (Alves et al., 2013), as well as for its regulation of fear expression (Gehrlach et al., 2019; Klein et al., 2021); vCA1 for its importance for CFC encoding (Kim & Cho, 2020) and recent recall (Jimenez et al., 2020); the BLA for the role of BLA to PL projections in memory encoding (Kitamura et al., 2017; Klavir et al., 2017) and PL to BLA projections in memory recall (Do-Monte et al., 2015; Kitamura et al., 2017); and the CLA as CLA to EC projections are necessary during memory encoding (Kitanishi & Matsuo, 2017) and for its involvement in attention (Atlan et al., 2018).

To assess the relative activity of these PL inputs during fear memory consolidation, we needed a tracing technique that could be coupled with neuronal activity measurements from encoding to remote recall, which cannot be achieved with rabies tracing from engram cells. Therefore, we combined conventional retrograde tracing with neuronal activity-dependent cFos staining: Injection of AAVretro-GFP in PL prior to any behavioral test allowed to trace all anatomical projections to the PL (Figure 3A, B), while cFos IHC 90min after CFC encoding, recent and remote memory recall allowed to assess the activation of these projections (Figure 3A). In each region we measured cFos as well as GFP traced inputs (Figure 3 S1, S2), thus controlling for homogenous tracing across behavioral groups. Next, we compared the pattern of activation between PL projectors only and the region as a whole to highlight the specific recruitment of PL projectors, and we focused on the associative information conveyed in this activity by normalizing it to the no shock control groups (Figure 3, see Figure 3 S1 and S2 for quantifications only normalized to chance level).

**Figure 3.**
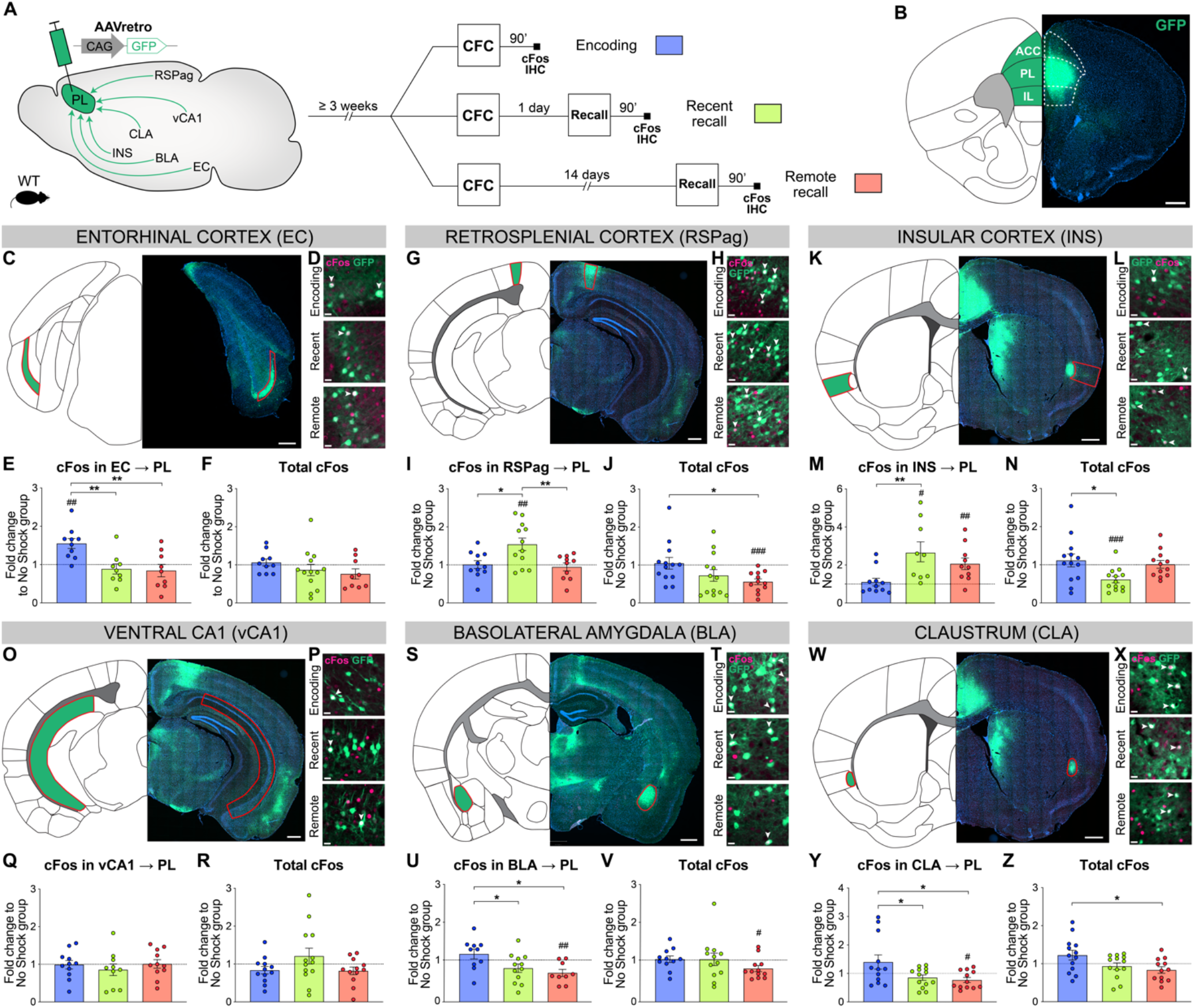
PL inputs are differentially activated during memory consolidation. **(A)** Experimental design, injection of AAVretro-GFP in the PL for input tracing, and quantification of activation by cFos immunostaining 3 weeks later at either CFC encoding (blue), recent (green) or remote (red) recall. Brains were collected for 90 minutes after the behavior session. **(B)** Representative image of AAVretro-GFP injection site in the PL region of the mPFC. Scale: 500μm. **(C-Z)** For each region: Representative image of PL input tracing, magnified view of GFP and cFos at encoding, recent and remote timepoints (all scales: 20μm); quantifications of cFos in PL projections and total cFos in the input region, expressed as fold change to the No Shock control group. Note that cFos in PL projections values were first normalized by chance level for each animal (see Figure 3 S1, S2). **(C-F)** EC (**C**, scale 500μm), **(E)** cFos in EC → PL (one way ANOVA, F (2, 25) = 8.153, p = 0.0019) and **(F)** total cFos. **(G-J)** RSPag (**G**, scale 400μm), **(I)** cFos in RSPag → PL (one-way ANOVA, F (2, 35) = 3.275, p = 0.0497) and **(J)** total cFos (one-way ANOVA, F (2, 35) = 3.275, p = 0.0497). **(K-N)** INS (**K**, scale 500μm), **(M)** cFos in INS → PL (one-way ANOVA, F (2, 27) = 5.405 p = 0.0106) and **(N)** total cFos (one-way ANOVA, F (2, 35) = 4.583, p = 0.0171). **(O-R)** vCA1 (**O**, scale 400μm), **(Q)** total cFos and **(R)** cFos in vCA1 → PL. **(S-V)** BLA (**S**, scale 500μm), **(U)** cFos in BLA → PL (one way ANOVA, F (2, 28)= 4.922, p = 0.0147) and **(V)** total cFos. **(W-Z)** CLA (**W**, scale 500μm), **(Y)** cFos in CLA → PL (one-way ANOVA, F (2, 34) = 4.502, p = 0.0184) and **(Z)** total cFos (oneway ANOVA, F (2, 35) = 3.833, p = 0.0313). Stars represent p-values of Tukey’s multiple comparisons tests (*: 0.01<p<0.05; **: 0.001<p<0.01), hashtag signs represent p-values of twotailed one sample t-tests comparing the difference to 1, which represents the No Shock controls (#: p≤0.05; ##: 0.001<p≤0.01; ###: p≤0.001). n = 9-13 animals per group.

We first investigated cortical areas projecting to PL: EC (specifically layer 5, comprising most of EC traced cells, Figure 3C), RSPag (which contained most of RSP traced cells, Figure 3G), and INS (Figure 3K). In the EC, we observed a significant activation of PL projections at encoding compared to both recent and remote recalls (Figure 3E), which was not the case in total cFos quantifications (Figure 3F). This suggests a specific recruitment of EC neurons projecting to PL (EC → PL) at encoding. In contrast, RSPag and INS displayed a different pattern of activation, as there was no activation in RSPag → PL and INS → PL projections at encoding, but during recent memory recall (Figure 3I and M, respectively). Compared to total cFos in both regions, this activity was again specific to PL projectors (Figure 3J and N, respectively).

Second, we investigated PL inputs in subcortical areas: vCA1 (Figure 3O), BLA (Figure 3S) and CLA (Figure 3W). In vCA1, we did not observe a differential recruitment of vCA1 → PL projections between different times of memory consolidation (Figure 3Q, R; Figure 3 S2D). However, the elevated cFos expression in the vCA1 as a whole at encoding as well as in the no shock group supports the role of HPC in context exploration (Figure 3 S2B) (Schiller et al., 2015). For the BLA, we observed a significant activation of BLA → PL projections at encoding compared to the recalls (Figure 3U), which is not the case for total cFos in BLA (Figure 3V). The recruitment of BLA → PL projection at encoding is in agreement with its importance during fear memory formation (Kitamura et al., 2017; Klavir et al., 2017). Interestingly, we observed the same pattern of activation in CLA → PL projections (Figure 3N), together with an overall higher activation of the whole CLA region at encoding compared to remote recall (Figure 3M).

Taken together, we found that PL inputs from the EC, BLA and CLA were active only at encoding, while RSPag and INS projections were recruited during recent memory recall.

### PL inputs are functionally relevant at different stages of memory consolidation

Next, in order to establish whether the differential activity in PL inputs across memory consolidation is also functionally relevant, we selectively inhibited each projection at the time(s) when they were most active and tested subsequent memory retention. We used the Designer Receptor Exclusively Activated by Designer Drug (DREADD) receptor hM4Di, which upon Clozapine-N-Oxide (CNO, the DREADD agonist) administration inhibits neuronal activity (Roth, 2016). We targeted hM4Di expression to specific PL inputs by injecting AAVretro-Cre into the PL, and AAV-DIO-hM4Di-mCherry (or AAV-DIO-mCherry for controls) in the desired input region (Figure 4A-F).

**Figure 4.**
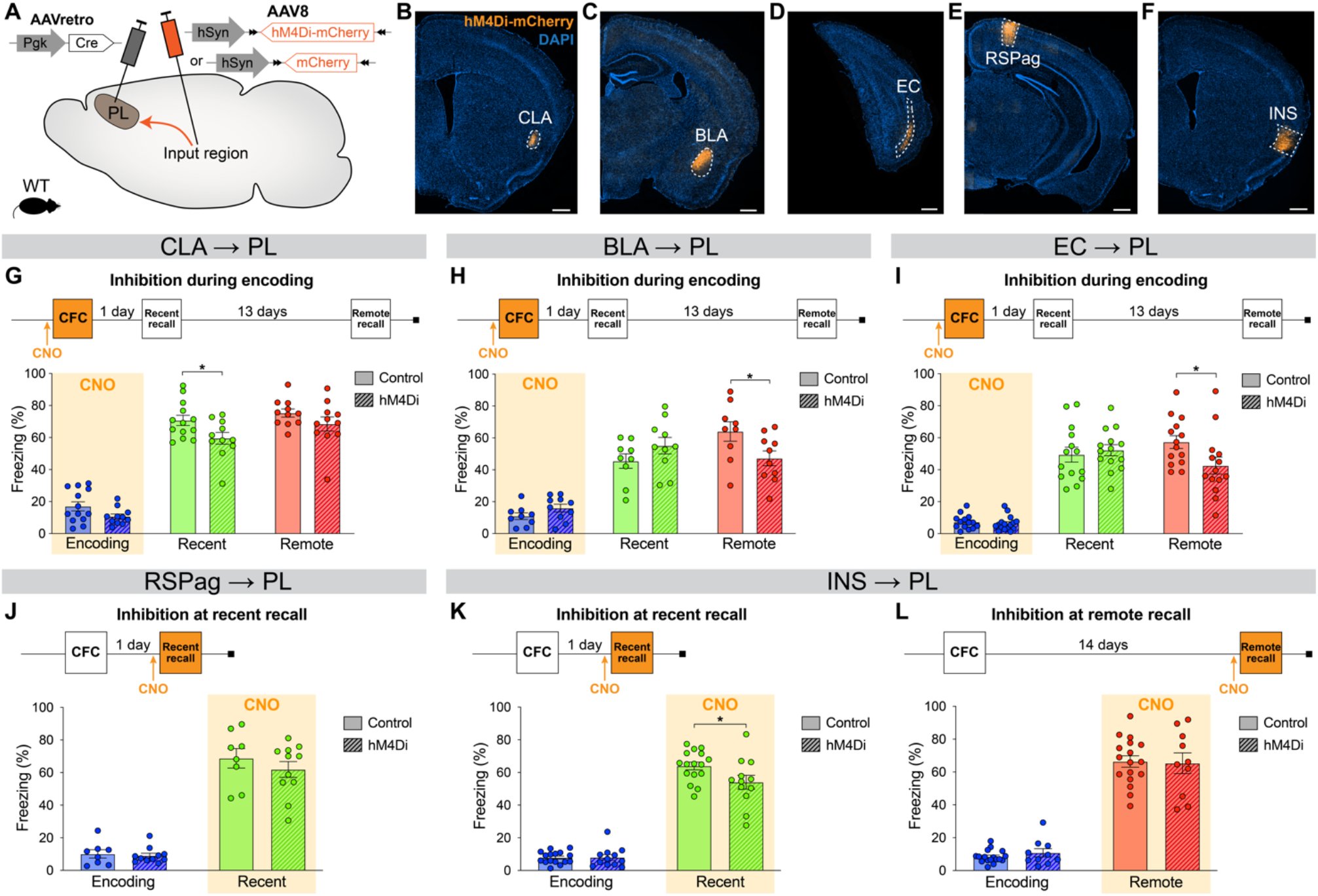
Chemogenetic manipulation of PL inputs reveals the functional importance of CLA projections at encoding and INS projections at recent memory recall. **(A)** Experimental design. AAVretro-Cre was injected in the PL, and AAV-DIO-hM4Di-mCherry (or AAV-DIO-mCherry for controls) in the desired input region in order to specifically inhibit the projections to the PL upon CNO injection. Representative images of the injection site in the input region for CLA **(B)**, BLA **(C)**, EC **(D)**, RSPag **(E)** and INS **(F)**, all scales 500μm. Experimental timeline and freezing percentage of **(G)** CLA → PL inhibition during encoding, **(H)** BLA → PL inhibition during encoding, **(I)** EC → PL inhibition during encoding, **(J)** RSPag → PL inhibition during recent recall, **(K-L)** INS → PL inhibition during recent **(K)** and remote **(L)** recall. Stars represent p-values of two-tailed unpaired t-tests between hM4Di and control groups (*: p≤0.05). n = 8-17 animals per group.

First, we assessed the functionality of projections that were active at encoding, namely the CLA, BLA, and EC. For the CLA → PL inhibition, we observed an impaired memory at recent recall (Figure 4G). To confirm that CNO indeed inhibits hM4Di-expressing neurons, we expressed it in CLA → PL neurons (Figure 4 S1A) and perfused the animals 90 minutes after CFC to stain for cFos (Figure 4 S1B, C). While the percentages of hM4Di^+^ and cFos^+^ cells were equivalent in both groups (Figure 4 S1D, E), the amount of double positive hM4Di^+^cFos^+^ cells was significantly decreased with CNO injection, confirming the inhibition of projection neurons during behavior (Figure 4 S1F). Furthermore, this behavioral result was not due to a delayed effect of CNO injection, as inhibiting CLA → PL projections right after encoding did not result in impaired memory at any time point (Figure 4 S2A, B). Likewise, the effect of CLA → PL inhibition was not due to an unspecific effect on locomotion and exploratory behavior as tested in an open-field arena (Figure 4 S2C-E). In contrast to the effect of CLA → PL inhibition, when BLA → PL and EC → PL projections were inhibited during encoding, we observed an impairment of remote memory recall for both, while recent recall was unaffected (Figure 4H and I, respectively). These results indicate that while the BLA → PL and EC → PL projections are necessary at encoding for the consolidation of remote memories, as shown previously (Kitamura et al., 2017), the CLA → PL projection is necessary at encoding for recalling recent memories.

Next, we tested the functionality of projections that were most active during fear memory recall, namely the RSPag and INS to PL projections. We found that although the RSPag → PL projection was specifically active at recent recall (Figure 3I), its inhibition during this time did not affect memory retrieval (Figure 4J). Interestingly, it was recently reported that although the whole RSP is necessary for recent and remote recall, it is rather the granular subregion of the RSP and not the RSPag that is responsible for this effect, suggesting a dissociated role of the two RSP subregions which could explain our observations (Tsai et al., 2022). In contrast, INS → PL projection inhibition during recent recall resulted in decreased freezing (Figure 4K). Lastly, consistent with no significant activation of the INS → PL projection at remote memory recall, the inhibition of this projection did not result in any behavioral effect (Figure 4L).

Taken together, these findings indicate that CLA, BLA and EC projections to the PL are necessary at encoding, but with different time implications. While the BLA and EC connections are important for recalling remote memories, the CLA projection is specifically important for recalling recent ones. In addition, recent memory recall is also under the influence of the INS → PL projection, since its inhibition at this time led to significant memory impairment. This suggests a progressive functional shift in PL projections regulating memory consolidation.

### PL engram reactivation correlates with memory retrieval when CLA or INS inputs are inhibited

Lastly, we decided to further investigate the effect of CLA and INS input inhibition on engram reactivation in the PL. We hypothesized that if the inhibition of a specific PL input results in memory impairment, then the reactivation of the original PL engram, established at the time of memory encoding, may also be altered. Indeed, engram reactivation has been correlated with memory retention in BLA (Reijmers et al., 2007), and artificial engram reactivation in the PL (Kitamura et al., 2017) or HPC (Liu et al., 2012) has been found to trigger memory recall. In order to measure engram reactivation, we used the cFos::tTA mouse line (Reijmers et al., 2007), expressing the Doxycycline (Dox)-dependent tTA transcription factor under the cFos promoter, which we injected with AAV-TRE-GFP into the PL 3 weeks before CFC (Figure 5A). As tTA specifically binds the TRE (tetracycline responsive element) promoter in absence of Dox, this approach allows for inducible long-term expression of GFP in PL engram cells during a desired time-window (Figure 5B, C). In combination with chemogenetic inhibition of projection neurons as previously described (Figure 4), we then silenced selective PL inputs and assessed the degree of engram reactivation between CFC encoding and recall, by measuring cFos and GFP overlap in PL (Figure 5C).

**Figure 5.**
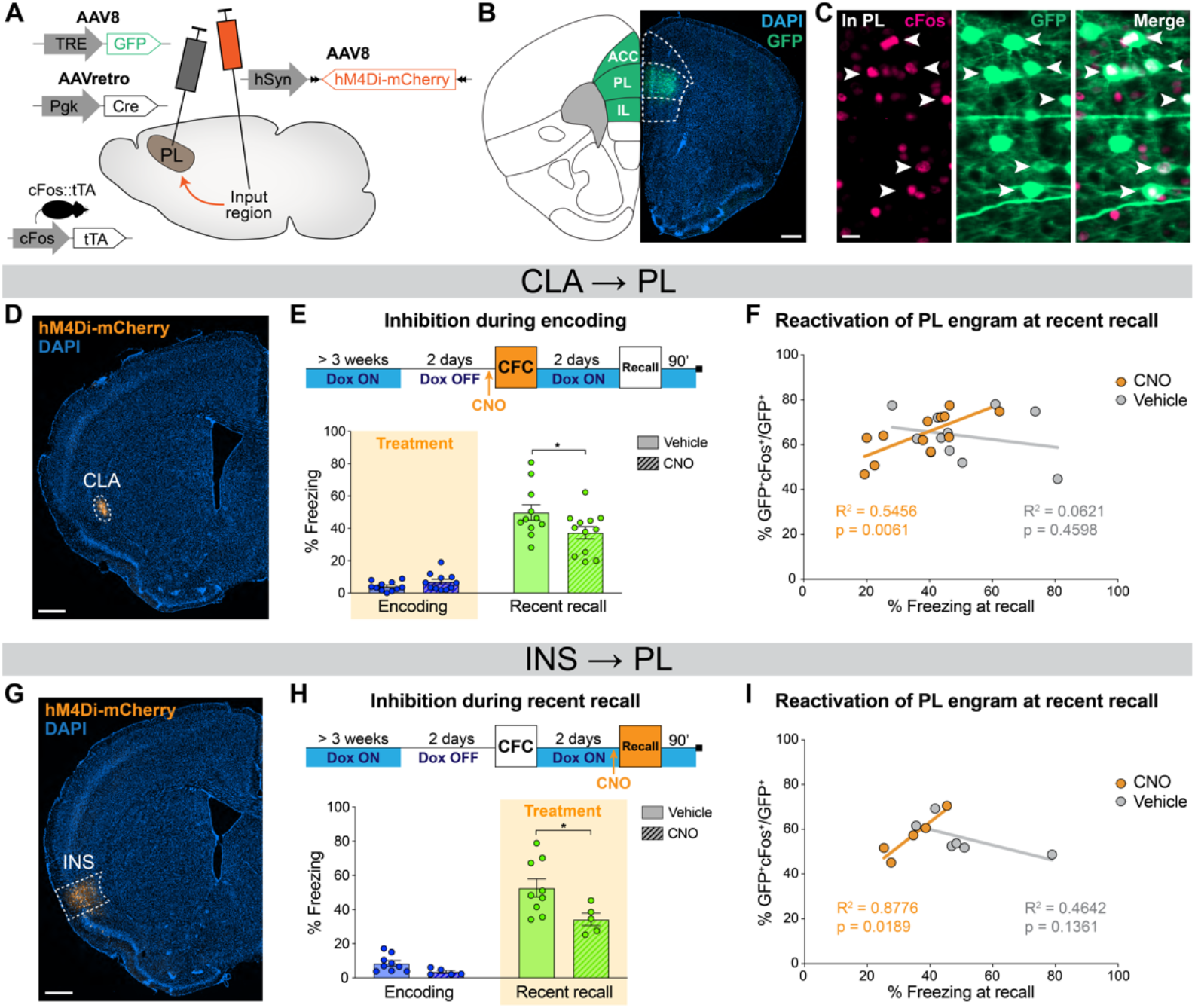
PL engram correlates with freezing when CLA or INS inputs are inhibited. **(A)** Experimental design. 3 weeks before behavior started, cFos::tTA mice were injected with AAVretro-Cre in PL and AAV-DIO-hM4Di-mCherry in the desired input region, as well as AAV-TRE-GFP in PL, so that GFP was only expressed in cFos^+^cells in absence of Doxycycline (Dox). **(B)** GFP expression at the PL injection site (scale 400μm). **(C)** Magnified view in PL (scale 20μm) of reactivated engram cells, indicated by white arrows. **(D)** Representative image of the CLA input region. **(E)** Experimental timeline (top) and freezing percentage (bottom) during recent memory recall when CLA → PL projections were inhibited during encoding. **(F)** Reactivation of PL engram cells (%GFP^+^cFos^+^/GFP^+^) at recent recall for CLA → PL inhibition, correlated with freezing percentage at recent recall for CNO (orange) and vehicle (grey) groups. **(G)** Representative image of the INS input region. **(H)** Experimental timeline (top) and freezing percentage (bottom) during recent memory recall when INS → PL projections were inhibited during recent recall. **(I)** Reactivation of PL engram cells (%GFP^+^cFos^+^/GFP^+^) at recent recall for INS → PL inhibition, correlated with freezing percentage at recent recall for CNO (orange) and vehicle (grey) groups. **(E, H)** Stars represent p-values of two-tailed unpaired t-tests between CNO and vehicle groups (*: p≤0.05). **(F, I)** Correlations assessed with linear regressions, R^2^ and p-values are reported on the graphs. n = 11-12 (CLA) or 5-9 (INS) per group.

First, we focused on the CLA → PL inhibition at memory encoding (Figure 5D). Behaviorally, this approach confirmed the impaired memory at recent memory recall as observed in wild-type mice (Figure 5E, see also Figure 4G). Furthermore, we found a significant correlation between PL engram cells reactivation, measured as double positive GFP^+^cFos^+^ cells normalized to the total number of GFP^+^ cells, and freezing at recent recall, which was observed only in the CNO group (Figure 5F). No differences were observed in overall GFP^+^, cFos^+^, double positive GFP^+^cFos^+^ and total reactivation percentages in PL between CNO- and vehicle-treated animals. (Figure 5 S1A-D). In contrast, when we inhibited the CLA → PL projection during encoding and tested remote recall, memory was not impaired (Figure 5 S2B, see also Figure 4G) and we observed no effect on PL engram reactivation (Figure 5 S2C-G). This result indicates that CLA → PL inhibition during encoding modifies PL engram reactivation at recent recall only, and that following this inhibition, the less animals reactivate the original PL engram, the less they recall the fear memory.

Second, we inhibited the INS → PL projection at recent recall (Figure 5G), which also confirmed the behavioral effect on memory retrieval (Figure 5H) in wild type mice (Figure 4K). Similar to the CLA results, there was no difference between the CNO and vehicle groups in the percentage of GFP^+^, cFos^+^, double positive GFP^+^/cFos^+^ cells or total reactivation (Figure 5 S1E-H). However, we again observed a significant correlation in the CNO group between PL engram reactivation and freezing at recent recall (Figure 5I), indicating that INS → PL inhibition at recent recall impairs recent memory retrieval and modifies PL engram reactivation.

These findings suggest that the spatiotemporal shift in the activity and functionality of PL projections during memory consolidation also occurs at the level of PL engram cells.

## Discussion

In this study, we investigated the role of specific PL inputs during the course of fear memory consolidation. Using an unbiased tracing approach combined with pathway-specific chemogenetic inhibition experiments, we discovered a novel functional implication of two PL inputs, namely from the CLA and INS, and confirmed the role of two others, from the BLA and EC. More precisely, we found that the BLA → PL and EC → PL projections are necessary at encoding for remote memory recall, confirming previous results (Kitamura et al., 2017), while the CLA → PL projection is necessary also at encoding, but specifically for recent memory recall. In addition, we found the INS → PL pathway to be necessary for memory expression during recent recall.

These results expand the existing literature on memory consolidation and refine the working model of memory formation and retrieval. Importantly, our data add to the growing evidence on the importance of the mPFC during early phases of memory consolidation (Bero et al., 2014; Cho et al., 2017; Rajasethupathy et al., 2015; Takehara-Nishiuchi et al., 2020; Zelikowsky et al., 2013). Thereby it further challenges the standard theory of memory consolidation, which posits that the HPC is necessary for encoding and recent recall, while the mPFC would take over from the HPC only at remote recall (Frankland & Bontempi, 2005). Indeed, we observe a significant activation of the PL during memory encoding already, as well as the necessity of several of its inputs at encoding and recent recall, which advances the temporal engagement of the mPFC to earlier than remote recall only.

At remote memory recall, in contrast, we did not observe any PL input that was engaged or behaviorally relevant. To our knowledge, such an input has never been reported, although the importance of mPFC as a whole (Bero et al., 2014; Frankland et al., 2004; Goshen et al., 2011), and of its functional outputs at remote recall is well established (Do-Monte et al., 2015; Kitamura et al., 2017). This apparent gap, or the impossibility to trace back the flow of information upstream of the mPFC, could be explained if the inputs are distributed across a vast network after memory consolidation. In that case, they could potentially be redundant and therefore harder to functionally identify. As such, these results are in agreement with the multiple trace theory (Moscovitch et al., 2006; Nadel & Moscovitch, 1997), which posits that the HPC first encodes memory upon learning, but does not store the memory per se. Rather, it is a distributed cortical network of neurons – with inputs from the HPC – that represents pieces for long-term information storage. With time, this leads to the creation of multiple traces in the brain for a given memory, making it more stable and less likely to be disrupted, as we have observed here.

Corroborating the implication of the PL during the early phases of memory consolidation, we found that the CLA → PL is necessary at encoding, which is the first report that this PL input is functionally important during fear memory formation. The CLA has known roles in attention (Atlan et al., 2018) and context exploration (Kitanishi & Matsuo, 2017), which are likely to support its role in memory formation. Interestingly, we observed that the CLA → PL projection was necessary at encoding only for recent recall, and not for a later remote recall. This result suggests that this projection has a time-limited effect on memory consolidation, and that other brain areas allow for a proper remote recall and thereby compensate in case of the CLA → PL inhibition at encoding. Of note, a CLA → EC projection has also been reported to be necessary at CFC encoding for recent recall (Kitanishi & Matsuo, 2017), while here we have observed that the EC → PL projection is functional at encoding for remote recall. This opens the possibility of an indirect circuit from the CLA to the PL via the EC, which could compensate in case the CLA → PL direct connection is impaired, but this remains to be experimentally determined. Nevertheless, these results imply a shift in PL circuits underlying contextual fear memory encoding, reminiscent of findings that reported a shift in PL circuits when an auditory fear memory is retrieved (Do-Monte et al., 2015).

In line with the notion of an overall shift in PL circuits during early memory consolidation, we report that the INS → PL projection is necessary at recent, but not at remote memory recall. The INS as a whole has classically been involved in taste learning (Yiannakas & Rosenblum, 2017) and the encoding of conditioned taste aversion (Sano et al., 2014), but has also been been implicated in recent CFC recall (Alves et al., 2013) as well as in auditory fear memory extinction (Klein et al., 2021). The INS to mPFC reciprocal connectivity has only been investigated in the context of taste learning, where it was recently found necessary for the expression of novel taste aversion (Kayyal et al., 2021). However, a direct role of INS input to PL during fear memory consolidation has not been described before. This finding therefore supports a broader role for the INS in learning beyond taste-related tasks (Boughter & Fletcher, 2021; Shi et al., 2020).

Since the CLA and INS manipulations both resulted in impaired recent memory recall, we decided to assess PL engram cells for their reactivation, a characteristic of engram cells that is linked with memory performance during recall (Kitamura et al., 2017; Liu et al., 2012; Reijmers et al., 2007). We found that the reactivation of PL engram cells significantly correlated with freezing behavior, but only when the CLA and INS inputs were inhibited, and not in controls. This finding raises the question of the functionality of PL engram cells at recent recall in normal conditions. Recently, a concept of “silent” engram cells in the PL has been developed, which postulates that silent PL engram cells have the particular feature of not being activated by recent recall – although their artificial reactivation can trigger recall at this time –, but of becoming active only at remote recall (Kitamura et al., 2017; Matos et al., 2019). Our findings are thus in line with this concept: When the CLA → PL projection is inhibited at encoding, or when the INS → PL projection is inhibited at recent recall, overall engram reactivation is left unchanged, but recent memory expression is impaired, which would suggest that the PL engram population itself is not required for recent memory recall. Since upon inhibition we observe the emergence of a correlation between PL engram cells reactivation and memory retention, it is possible that CLA and INS inputs target inhibitory neurons in the PL, which normally prevent engram reactivation during recent recall, thus allowing the PL engram cells to stay functionally silent. Releasing this inhibition could perturb normal memory expression, explaining the observed memory impairment at recent recall. Indeed, it has been reported that CLA → mPFC targets inhibitory neurons (Jackson et al., 2018), and that PL interneurons are necessary for memory encoding (Cummings & Clem, 2020), but this hypothesis remains to be tested at the level of PL engram cells.

There are several experimental limitations that accompany these findings. First, the use of two different engram-tagging mouse lines, TRAP2 for the rabies tracing and cFos::tTA for the engram reactivation experiments, was dictated by technical constraints. A TRE-dependent rabies tracing system was not readily available at the start of this study, and the use of a Cre-dependent system for chemogenetic inhibition precluded the use of the TRAP2 line again for the engram reactivation experiments. However, as these two mouse lines are both cFos promoter-based (DeNardo et al., 2019; Reijmers et al., 2007), we would not expect major differences with the use of one or the other lines. Indeed, rabies tracing from PL engram cells using the cFos::tTA line has been published since, and the reported input areas are all also found in our brain-wide screen (Kitamura et al., 2017). Second, the use of two different retrograde tracing viruses raises the question of tropism. Indeed, RVΔG and AAVretro have been reported to not trace the exact same set of input regions, notably with AAVretro being biased towards cortical inputs (Sun et al., 2019). We could have therefore missed some regions due to preferential input tracing. A third limitation are the relatively slow kinetics of chemogenetic inhibition. As CNO is injected 30 minutes before behavior, we cannot exclude that compensation mechanisms may take over, especially in the case of remote recall inhibition, which would prevent a functional isolation of the targeted projection during behavior as reported previously (Goshen et al., 2011). By restricting the inhibition to the smallest possible period, the use of optogenetics could allow to visualize the consequences of this inhibition in real time in future experiments. Lastly, differences in timing and strength of behavioral protocols could explain discrepancies with other studies. For example, using cFos IHC, we did not observe an increased activity at remote recall in mPFC regions as opposed to previous findings (Frankland et al., 2004). Unified conditioning protocols could help to clarify these.

These limitations notwithstanding, here we have shown that PL circuits undergo a spatiotemporal shift during contextual fear memory consolidation, with claustral inputs being critical at encoding, and insular cortical inputs at recent memory recall. Our results therefore support a dynamic and distributed nature of memory formation and storage.

## Materials and methods

### Animals

All animals and procedures used in this study were approved by the Veterinary Office of the Federal Council of Switzerland under the animal experimentation licenses VD2808.1 and VD2808.2. C57Bl/6JR wild-type male mice were purchased from Janvier Labs (France) around 6-7 weeks old and left for at least one week before the beginning of the experiments. cFos::Cre^ERT2^ (TRAP2) animals were bred in house from the original Jax strain #030323 on a C57Bl/6JR background. cFos::tTA male mice were bred in house from the original JAX strain #018306 on a C57Bl/6JR background. Animals were housed in a 12h light/dark cycle with water and food available ad libitum. All animals were group-housed except for the input tracing experiment where they were single caged 2 days before the end of the experiment. They were all handled by the experimenter for at least 3 days before the first behavioral procedure to reduce stress levels.

All behavioral procedures were performed between 1pm and 5pm local time and animals were randomly assigned to experimental groups.

### Viral stereotaxic injections

#### Procedure

At 7-8 weeks, animals were anesthetized with a mix of Fentanyl (0.05 mg/kg), Midazolam (5 mg/kg) and Metedomidin (0.5 mg/kg), i.p. After shaving and subcutaneous injection of a local anaesthetic mix (Lidocaine 6 mg/kg and Bupivacaine 2.5 mg/kg), the animal was placed on a stereotaxic frame. The skin was disinfected with Betadine and opened with a scalpel. The skull was thoroughly cleaned, the orientation of the head was adjusted, and holes were drilled at the desired coordinates with a 0.5mm drill bit. The virus was loaded into pulled glass capillaries (intraMARK, Blaubrand, tip diameter 10-20 μm), and injected to the target area at a speed of 100 nL/min. The needle was left in place for 5 min, and slowly pulled up to limit backflow. After all injections were done, the skin was sutured (5/0 Prolene, Ethicon), the animal was injected i.p. with Atipamezol (2.5 mg/kg) to reverse the anaesthesia, and placed back in a heated cage. After surgery, the animals were administered paracetamol in the drinking water for a week (Dafalgan, 1mg/mL).

#### Viruses

The following viruses were used in this study:

- AAV-DIO-TVA-2A-oG (Salk Institute Vector Core, serotype 8), here referred to as AAV-DIO-TVA-oG. Titer: 8.78×10^12^ GC/mL, mixed 1:1 with AAV-FLEX-GFP-oG (see below).
- AAV-EF1a-FLEX-H2B-GFP-P2A-oG (Salk Institute Vector Core, serotype 8), here referred to as AAV-FLEX-GFP-oG. Titer: 3.93×10^12^ GC/mL, mixed 1:1 with AAV-DIO-TVA-oG; total injection volume in PL: 400nL.
- Modified rabies virus RVΔG-mCherry, EnvA pseudotyped (Salk Institute Vector Core, SADB19 strain). Titer: 3.5×10^8^ ifu/mL. Injection volume in PL: 300nL.
- AAV-CAG-GFP (Addgene 37825, retrograde serotype), here referred to as AAVretro-GFP. Titer: 7×10^12^ GC/mL; injection volume in PL: 200nL.
- AAV-pgk-Cre (Addgene 24593, retrograde serotype), here referred to as AAVretro-Cre. Titer: 1.7×10^13^ GC/mL; injection volume in PL: 200nL.
- AAV-hSyn-DIO-hM4D(Gi)-mCherry (Addgene 44362 or Zürich VVF v84, serotype 8), here referred to as AAV-DIO-hM4Di-mCherry. Titer: 1.8×10^13^ GC/mL (Addgene – diluted ½) or 4.5×10^12^ GC/mL (VVF); injection volume: 150-200nL depending on the regions.
- AAV-hSyn-DIO-mCherry (Addgene 50459, serotype 8), here referred to as AAV-DIO-mCherry. Titer: 2.3×10^13^ GC/mL, diluted ½; injection volume: 150-200nL depending on the regions.
- AAV-TRE3G-GFP (UNC Vector Core, serotype 8), here referred to as AAV-TRE-GFP. Titer: 4.1×10^12^ GC/mL, mixed 1:1 with AAVretro-Cre; injection volume in PL: 250nL.

Injection coordinates from bregma:

1. - PL: AP +2.0; ML ±0.35; DV −2.2.
2. - EC: AP −4.15; ML ±3.55; DV −4.3.
3. - RSPag: AP −2.6; ML ±1.1; DV −0.6.
4. - INS: AP +1.0; ML ±3.85; DV −4.0 in WT mice, or AP +1.0; ML ±3.9; DV −4.0 in cFos::tTA mice.
5. - BLA: AP −1.0; ML ±3.15; DV −4.55.
6. - CLA: AP +1.0; ML ±3.2; DV −4.0 in WT mice, or AP +1.0; ML ±3.1; DV −4.0 in cFos::tTA mice.

For input tracing experiment with AAVretro-GFP, animals were injected unilaterally. For all other experiments, animals were injected bilaterally.

### Behavioral procedures

#### Contextual fear conditioning (CFC)

CFC encoding and recall were performed in the same chamber (TSE Systems). CFC encoding consisted in a first 3min exploration phase, followed by three 2s long 0.8mA foot-shocks spaced by 28s. After the last shock, the animal was left in the chamber for an additional 15s and brought back to its home cage. The recall consisted in a 3min exposure to the same context, without any shock. For all experiments except the engram reactivation, recent recall took place 1 day after the encoding and remote recall 14 days later. For the engram reactivation experiment, recent recall took place 2 days after the encoding, to leave enough time for GFP expression. The movement of the animals was automatically measured using an infrared beam cut detection system (TSE Systems). Freezing detection threshold was set at 1s of immobility. No shock control animals underwent the same procedure but did not receive any shocks. Animals without any chemogenetic manipulation were excluded if they froze less than 20% of the time during the recall (in total 2 animals were excluded in all experiments).

#### Tamoxifen injection

In the rabies tracing experiment, TRAP2 mice were injected i.p. with Tamoxifen (4-hydroxytamoxifen, Sigma-Aldrich, CAS 68392-35-8, 50 mg/kg) immediately after CFC. Tamoxifen was prepared as follows: powdered tamoxifen was dissolved in Ethanol 100% at a concentration of 20 mg/mL and stored at −20°C. On the day of the experiment, tamoxifen was re-dissolved by shaking at 37°C, 2 volumes of corn oil were added and ethanol was evaporated shaking at 37°C, for a final concentration at 10 mg/mL. Tamoxifen was kept at 37°C until injection to prevent precipitation.

#### Chemogenetic inhibition

In these experiments, mice were injected i.p. with Clozapine-N-oxide (CNO, Sigma-Aldrich, CAS 34233-69-7, 3 mg/kg) 30min before the desired behavioral phase. CNO was prepared as follows: 5mg of CNO were resuspended in 50μL of DMSO and stored at −20°C. On the day of the experiment, CNO was diluted 1/500 in NaCl to reach a concentration of 0.2 mg/mL, and injected at the desired volume. In the engram reactivation experiment, control animals were injected with an equivalent volume of vehicle i.p. (0.9% NaCl, B.Braun and 1/500 DMSO).

#### Open field test

For CNO control experiments, 30min after CNO injection the animals were placed in a large circular arena and left to freely explore for 15min. Video-tracking of the animals and locomotion quantification was automatically performed using the EthoVision software (Noldus).

#### Engram reactivation

cFos::tTA mice were administered Doxycycline (Dox, Sigma-Aldrich, CAS 24390-14-5) in the drinking water at 0.2 mg/mL. Dox was prepared as follows: Powdered Dox was resuspended in water from the animal facility at 50mg/mL, aliquoted and frozen at −20°C until further use. It was then diluted in water bottles to reach a concentration of 0.2 mg/mL. Dox was administered at least 3 weeks before the behavioral protocol started, and was refreshed every week. In order to open the tagging-window, Dox was removed 2 days before encoding, and administered back right thereafter for the remaining time of the protocol.

#### Sample size and behavioral replicates

No statistical method was used to predetermine sample size. The number of animals used in each experiment was the minimum required to obtain statistical significance, based on our experience with this behavioral paradigm and in agreement with standard literature. Data from the input tracing experiment was pooled from 3 independent batches (Figure 1 and 3 and their supplements). Data from the rabies tracing experiment comes from one batch (Figure 2 and its supplement). Data from the chemogenetic manipulation in WT was pooled from at least 2 independent batches for each manipulation (Figure 4). Data from the CNO controls comes from one batch each (Figure 4 S1 and S2 C-F). Data from the engram reactivation was pooled from 1-2 batches (Figure 5 and its supplements). In all graphs, one dot represents one animal.

### Histology

90min after the last behavioral test, animals were anesthetized with pentobarbital (150 mg/kg, Streuli Pharma) and transcardially perfused with first 1X PBS and then 4% paraformaldehyde (PFA) in 1X PBS. Brains were extracted, post-fixed overnight in 4% PFA, transferred in cryoprotectant (30% sucrose in 1X PBS) for at least 48h, and frozen at −80°C. Sections of 20μm were cut using a cryostat and kept free-floating in an antifreeze solution (30% ethylene glycol, 15% sucrose, 0.02% azide in 1X PBS) until staining.

For cFos immunostaining, sections were incubated in blocking buffer (1% BSA, 0.3% Triton-X in 1X PBS) for 90min at room temperature, followed by primary antibody incubation for 2 nights at 4°C in antibody dilution buffer (1% BSA, 0.1% Triton-X in 1X PBS). After 4 washes in 1X PBS + 0.1%Triton-X, they were incubated with secondary antibody in antibody dilution buffer for 2h at room temperature, stained with Hoechst (1:10.000 in 1X PBS, Invitrogen #H3570) for 5min and washed 3 times before mounting on glass slides and covered with Fluoromount-G mounting medium (Southern Biotech). Images were acquired on an Olympus slide scanner VS120 L100 with a 20X objective.

For the input tracing experiment, a primary antibody goat anti-cFos (1:1000, Santa Cruz #sc-52-G, RRID: AB_2629503) with a secondary antibody donkey anti-goat AF-647 (1:1000, Thermo Fisher Scientific #A21447, RRID: AB_2535864) was used. For all other experiments, a primary antibody rabbit anti-cFos (1:1000, Synaptic Systems #226003, RRID: AB_2231974) with a secondary antibody donkey anti-rabbit AF-647 (1:1000, Thermo Fisher Scientific #A31573, RRID: AB_2536183) was used. GFP and mCherry signals were not amplified.

For verification of the injection sites, 6 sections per animal were taken spanning the injection site, stained with Hoechst and mounted. Images were acquired on an Olympus slide scanner VS120 L100 with a 10X objective.

For the rabies tracing experiment, 1 every 4 sections of 20μm were mounted on Superfrost slides (Fisher scientific) and stained with Hoechst, before imaging on an Olympus slide scanner VS120 L100 with a 10X objective.

For simplicity and clarity in the text, we used “DAPI” to refer to nuclei stained with Hoechst.

### Image analysis

Images were analysed using QuPath (v0.1.4 to v0.3.1) (Bankhead et al., 2017), by an experimenter blinded to the groups.

For the rabies tracing experiment (Figure 2), every section was aligned to the reference Allen Brain Atlas using a Fiji plugin developed by the bioimaging platform at EPFL (Chiaruttini et al., 2022), before using a QuPath custom-built script for cell detection and classification (see supplementary material). It used multiple rounds of the built-in “Cell Detection” plug-in (once for each channel, plus one for DAPI). After detection, cells are given a classification based on the overlap of their coordinates to the DAPI channel detections.

For the input tracing experiment (Figure 3), 2-3 sections per brain region per animal were manually annotated based on the Allen Brain Reference Atlas, and identification of the detected GFP^+^and cFos^+^ cells within each annotated region was established using the custom-made QuPath script. An animal was excluded from further analysis if the percentage of traced inputs in a given area was below a region-specific threshold, as the amount of traced cells was region-dependent (thresholds were EC: 2%; RSPag: 1%; INS: 0.5%; vCA1: 0.5%; BLA: 0.6%; CLA: 1%). The chance ratio was calculated as (GFP^+^cFos^+^/DAPI)/chance level, where chance level was calculated as (GFP^+^/DAPI)x(cFos^+^/DAPI). Then, chance ratios were further normalized by the averaged chance ratio of the matching No Shock control groups (Figure 3). cFos^+^ cells in mPFC (Figure 1) were quantified in the non-injected contralateral mPFC using 3-4 sections per animal.

For the chemogenetic manipulation experiments (Figure 4-5), animals were excluded if the hM4Di-mCherry signal was leaking outside of the target region or if the amount of infected cells was too low. cFos^+^ and hM4Di^+^ in CLA (Figure 4 S1) were quantified using the QuPath custom-built script, in 3-4 sections per animal.

For engram reactivation experiments (Figure 5), cFos^+^ and GFP^+^ cells were quantified in PL using the QuPath custom-built script, in 3-4 sections per animal. Animals were excluded from further analysis if the percentage of GFP was below 1%.

### Statistics

All statistics and graphical representations were done with GraphPad Prism 9. All data are represented in mean ±SEM, with one dot representing one animal in all graphs. Data from the input tracing screen were analyzed using ordinary one-way ANOVAs, and further comparisons were performed with Tukey’s multiple comparisons tests between all groups (alpha = 0.05). In case of normalizations, difference to 1 was analyzed using two-tailed one sample t-tests (alpha = 0.05). Data from the chemogenetic manipulation and engram reactivation experiments were analyzed using two-tailed unpaired t-tests between the two groups (alpha = 0.05), and correlations were assessed with simple linear regressions.

## Author contributions

LD and JG conceptualized the project and wrote the paper; LD performed all experiments.

## Acknowledgements

We would like to thank all past and current members of the Laboratory of Neuroepigenetics for their support and discussion throughout this project, in particular Verena Doblmayr for her contribution to the behavioral experiments and histological analysis, Liliane Glauser for overall technical assistance and mice genotyping, and Gabriel Berdugo-Vega for insightful discussions on the project. We would like to also thank the EPFL Bioimaging platform (BIOP) for their continuous support in image acquisition and analysis, and the EPFL Center of Phenogenomics (CPG) for ensuring mice welfare at all times.

---

**Figure 2 – figure supplement 1.**
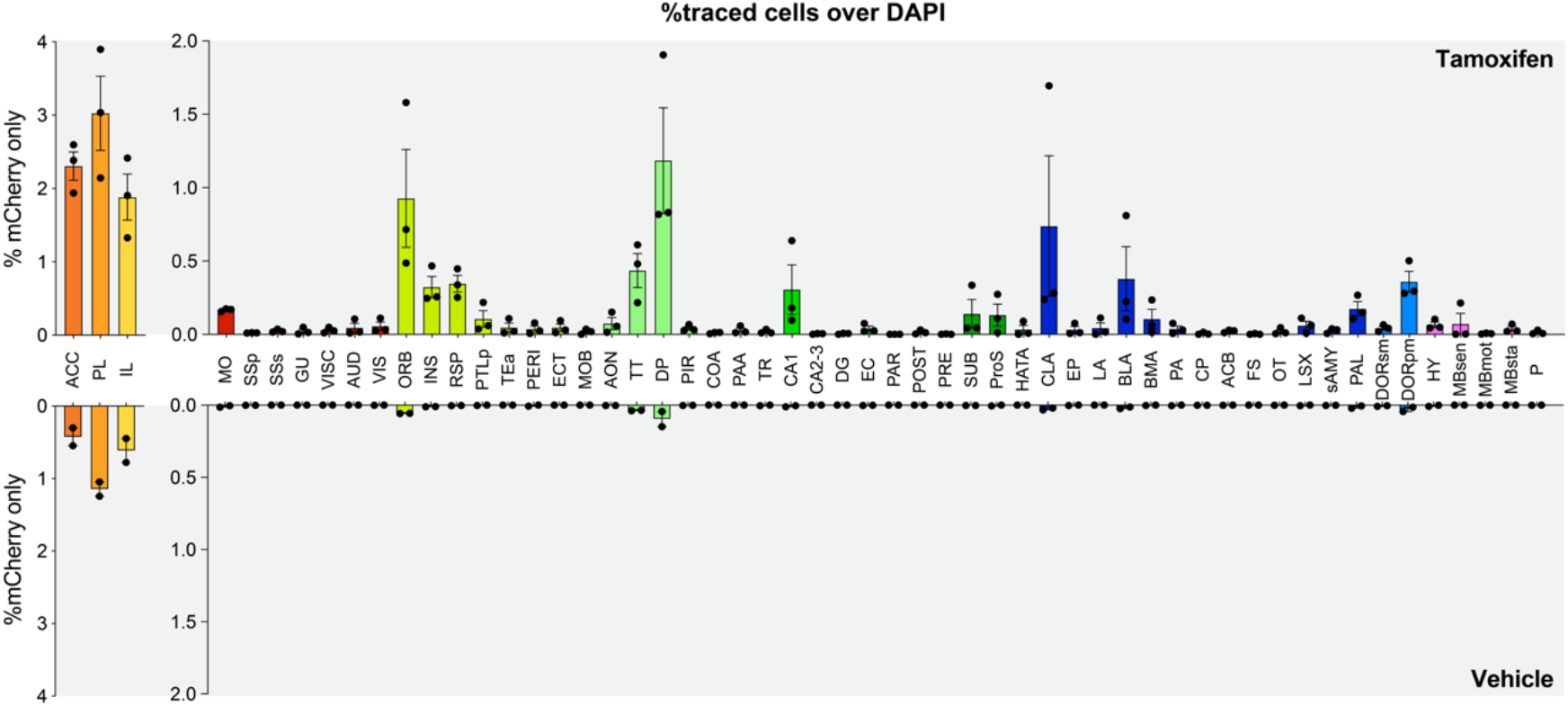
Raw quantifications of the rabies tracing experiment. Percentage of traced cells (mCherry^+^ only) in all regions with tamoxifen (top, n = 3 animals) or vehicle (bottom, n = 2 animals).

**Figure 3 – figure supplement 1.**
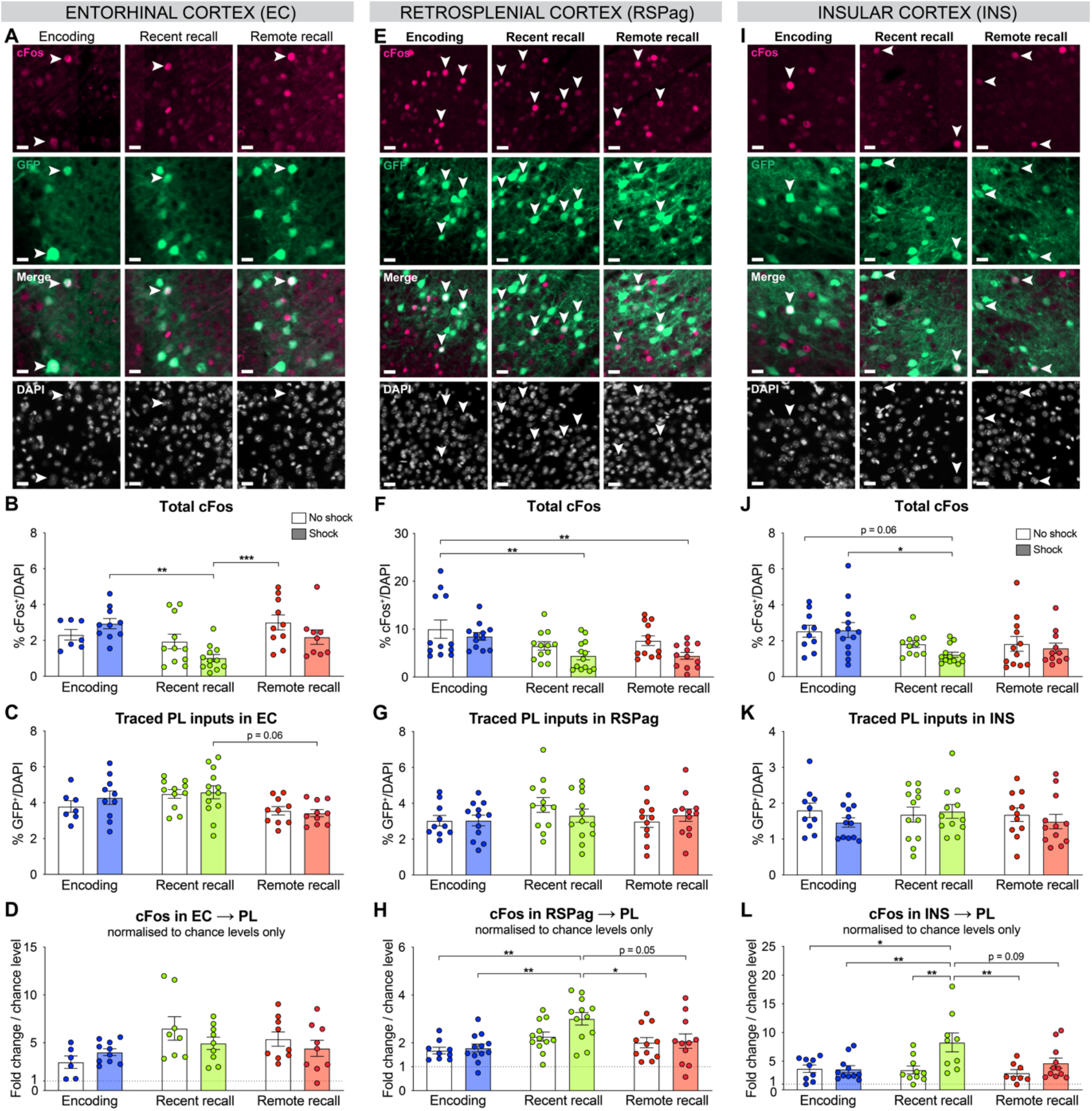
Complementary quantifications for the activation of PL inputs in EC, RSPag and INS. All representative images (from Figure 2) for EC **(A)**, RSPag **(E)** and INS **(I)**, scale 20μm. Total cFos percentage in EC (**B**, one-way ANOVA, F (5, 53) = 5.075, p = 0.0007), RSPag (**F**, one-way ANOVA, F (5, 67) = 4.156, p = 0.0024) and INS (**J**, one-way ANOVA, F (5, 64) = 2.971, p = 0.0179). Distribution of traced PL inputs across behavioral groups in EC (**C**, one-way ANOVA, F (5, 55) = 2.794, p =0.0255), RSPag **(G)** and INS **(K)**. Double positives (cFos^+^GFP^+^) normalized to chance levels in EC **(D)**, RSPag (**H**, one-way ANOVA, F (5, 61) = 4.618, p = 0.0012) and INS (**L**, one-way ANOVA, F (5, 52) = 4.312, p = 0.0023). Stars represent p-values of Tukey’s multiple comparisons tests (*:p≤0.05; **: 0.001<p≤0.01, ***: 0.0001<p≤0.001, ****: p≤0.0001). n = 9-13 animals per group.

**Figure 3 – figure supplement 2.**
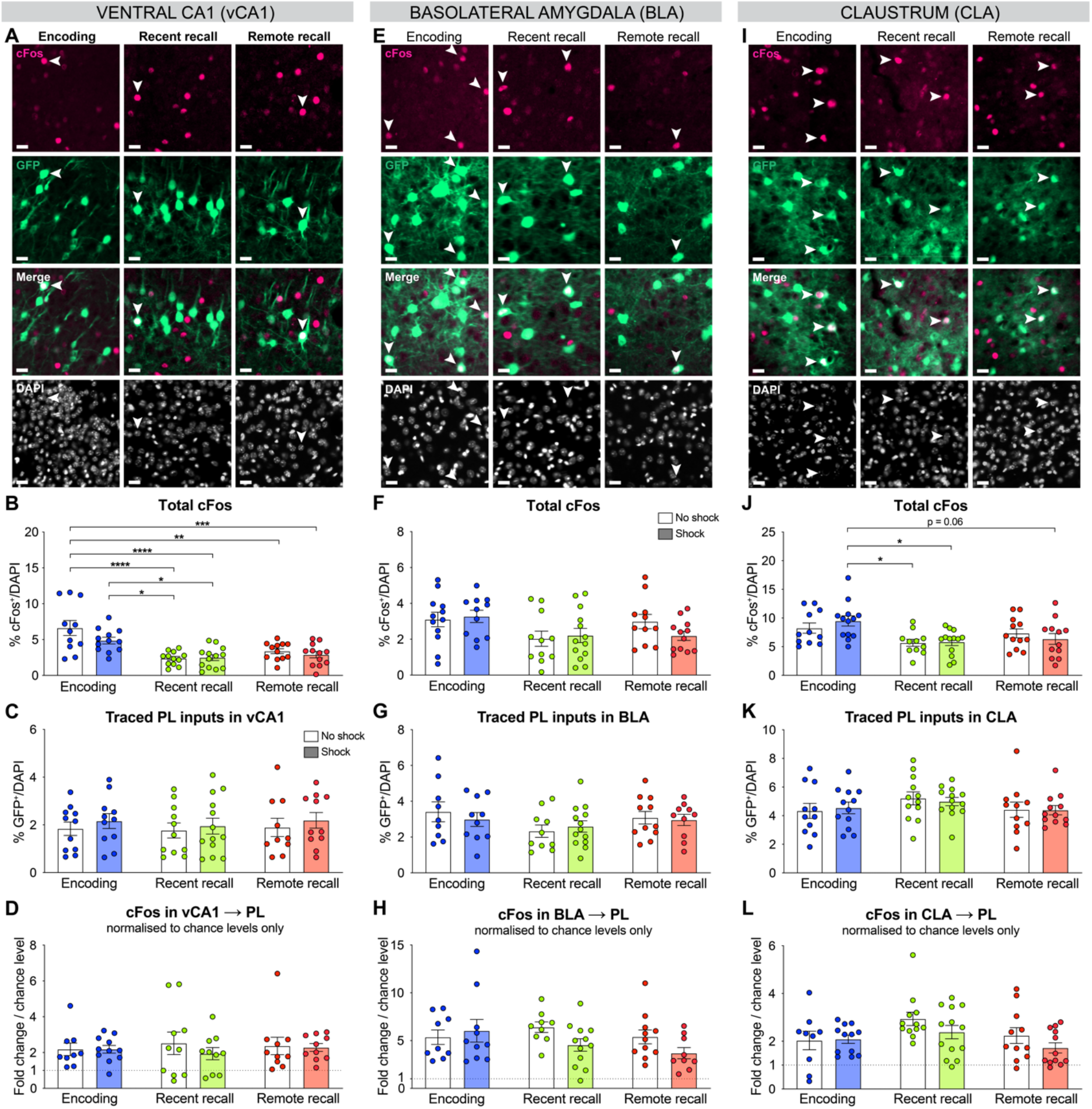
Complementary quantifications for the activation of PL inputs in vCA1, BLA and CLA. All representative images (from Figure 2) for vCA1 **(A)**, BLA **(E)** and CLA **(I)**, scale 20μm. Total cFos percentage in vCA1 (**B**, one-way ANOVA, F (5, 66) = 9.298, p < 0.0001), BLA **(F)** and CLA (**J**, one-way ANOVA, F (5, 66) = 3.593, p = 0.0062). Distribution of traced PL inputs across behavioral groups in vCA1 **(C)**, BLA **(G)** and CLA **(K)**. Double positives (cFos^+^GFP^+^) normalized to chance levels only in vCA1 **(L)**, BLA **(H)** and CLA (**L**, one-way ANOVA, F (5, 63) = 2.282, p = 0.0570). Stars represent p-values of Tukey’s multiple comparisons tests (*:p≤0.05; **: 0.001<p≤0.01, ***: 0.0001<p≤0.001, ****: p≤0.0001). n = 9-13 animals per group.

**Figure 4 – figure supplement 1.**
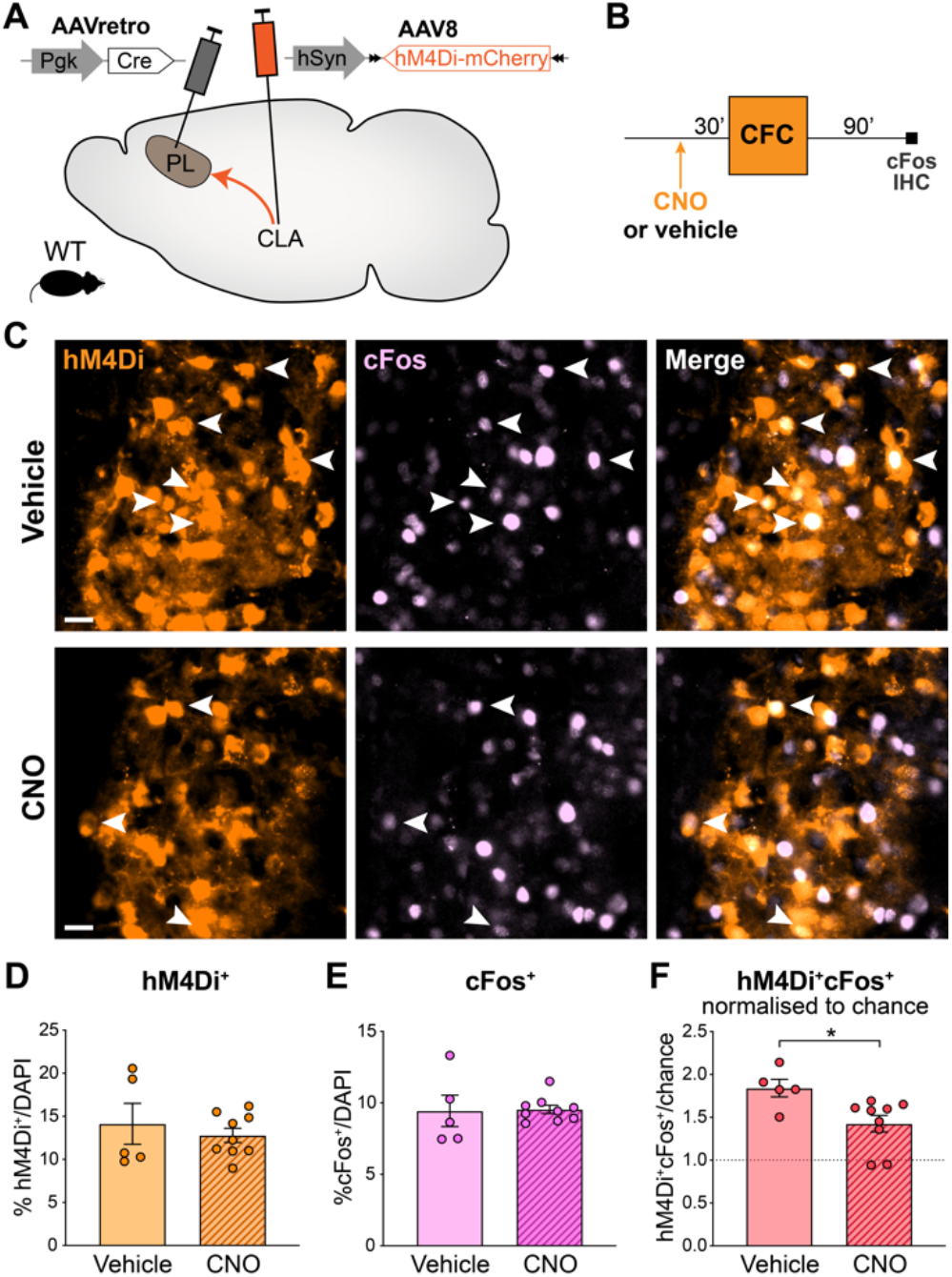
Quantification of the chemogenetic inactivation of CLA → PL projections. **(A)** Experimental design: AAVretro-Cre was injected in the PL and AAV-DIO-hM4Di-mCherry in the CLA, to express hM4Di specifically in CLA → PL projections. **(B)** Timeline of the experiment. CNO was injected 30 minutes before CFC, and mice were perfused 90 minutes later for cFos IHC. **(C)** Representative images of CNO and vehicle-injected groups, in the CLA. hM4Di in orange, cFos in light pink. Arrows indicate double positive hM4Di+cFos^+^cells. Scale 20μm. **(D)** Quantification of hM4Di^+^in CLA. **(E)** Quantification of cFos^+^in CLA. **(F)** Quantification of double positive hM4Di^+^cFos^+^cells in CLA, normalized to chance. Star represents p-value of two-tailed unpaired t-test between CNO and vehicle groups (*: p≤0.05). Vehicle: n = 5 animals; CNO: n = 9 animals.

**Figure 4 – figure supplement 2.**
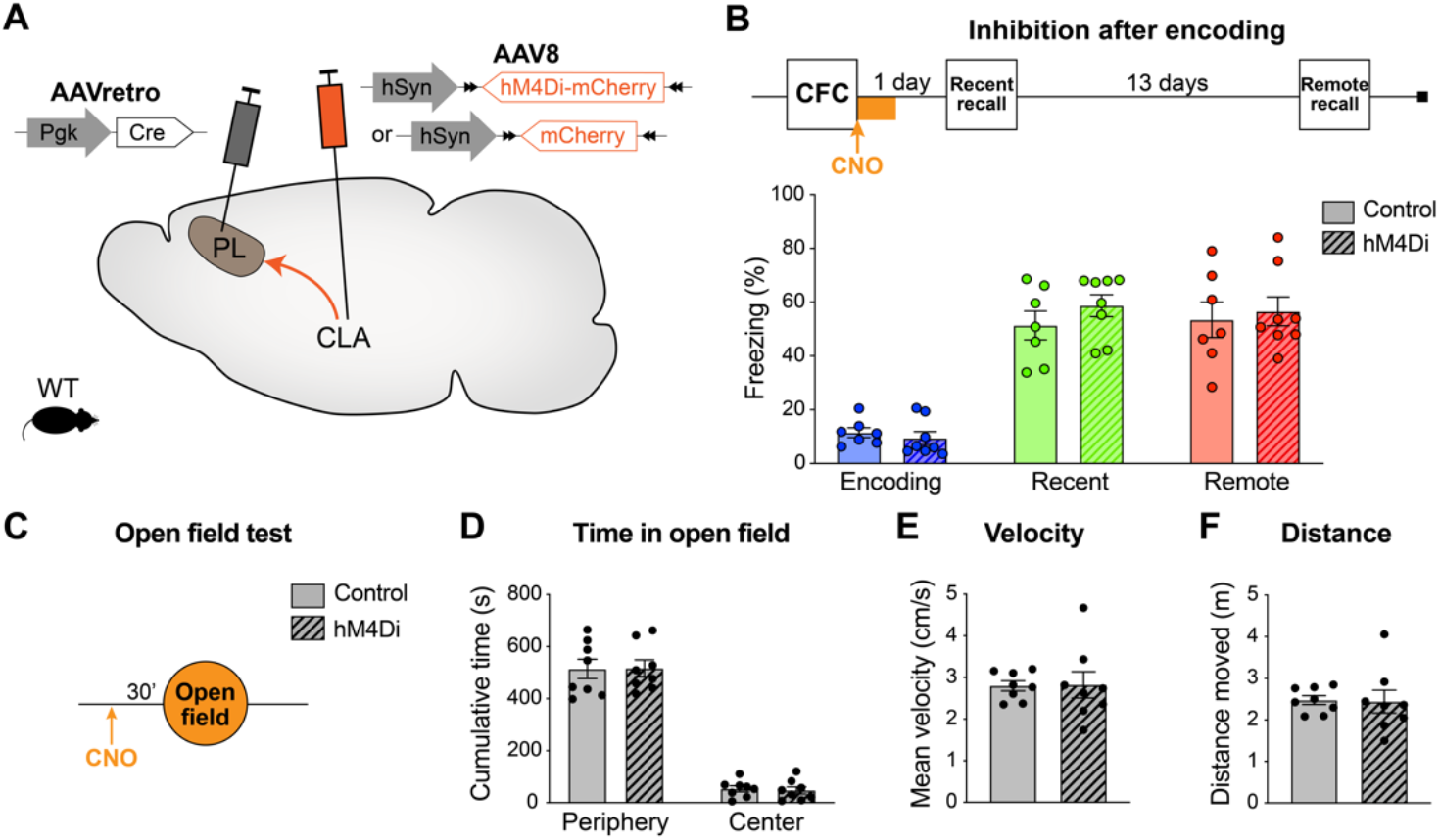
CLA → PL inhibition after encoding does not impair memory recall and does not alter locomotion and exploration behavior. **(A)** Experimental design: AAVretro-Cre was injected in the PL and AAV-DIO-hM4Di-mCherry in the CLA (or AAV-DIO-mCherry for controls), to express hM4Di (or mCherry) specifically in CLA → PL projection. **(B)** Experimental timeline and freezing percentage for CLA → PL inhibition after encoding. CNO was injected i.p. after CFC, and every 2 h, for a total of 4 injections. n = 7-8 per group. **(C)** Experimental timeline of the open field test, 30 minutes after a single CNO injection, measuring **(D)** time spent in the periphery or center of the arena, **(E)** velocity and **(F)** total distance moved. n = 8 per group.

**Figure 5 – figure supplement 1.**
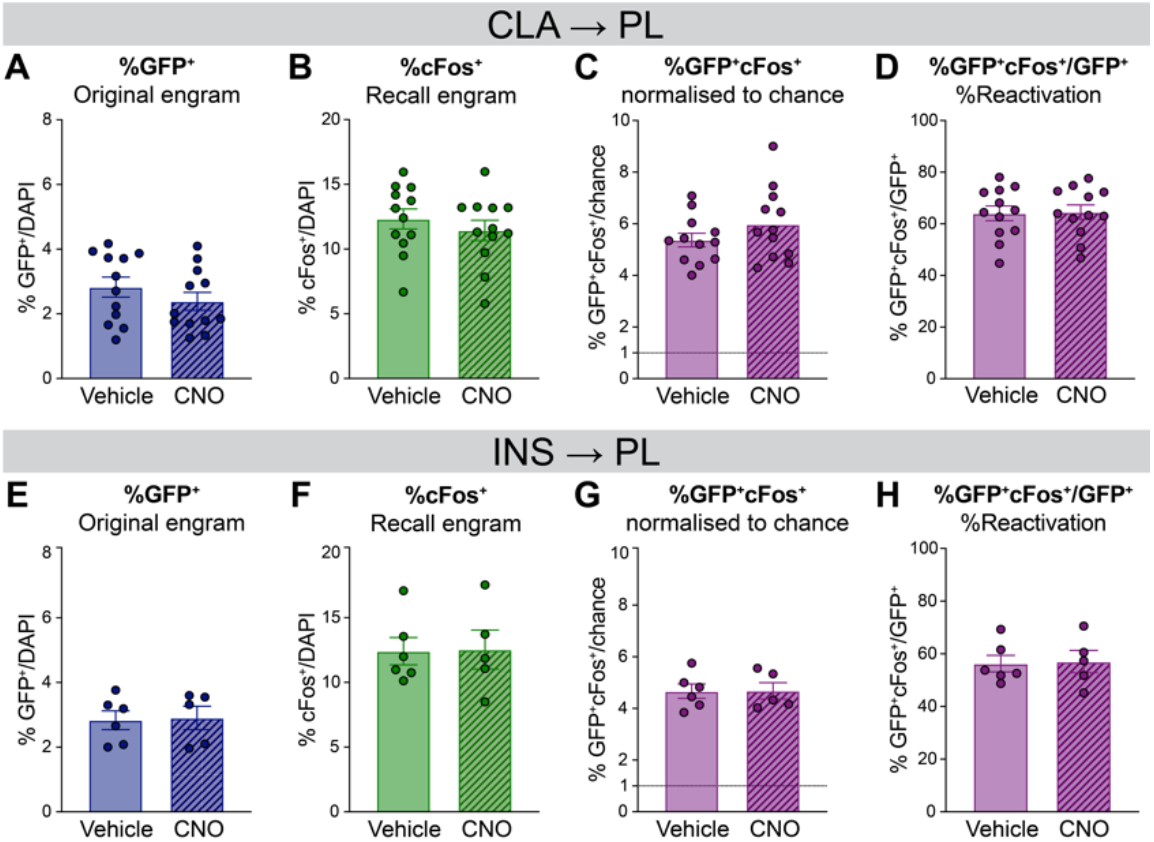
Complementary quantifications for the engram reactivation analysis in the PL. **(A-D)** For CLA → PL inhibition during encoding, percentage of **(A)** GFP^+^cells, **(B)** cFos^+^cells, **(C)** double positive GFP^+^cFos^+^cells normalized to chance and **(D)** reactivation expressed as %GFP^+^cFos^+^/GFP^+^; **(E-H)** For INS → PL inhibition during recent recall, percentage of **(E)** GFP^+^cells, **(F)** cFos^+^cells, **(G)** double positive GFP^+^/cFos^+^cells normalized to chance and **(H)** reactivation.

**Figure 5 – figure supplement 2.**
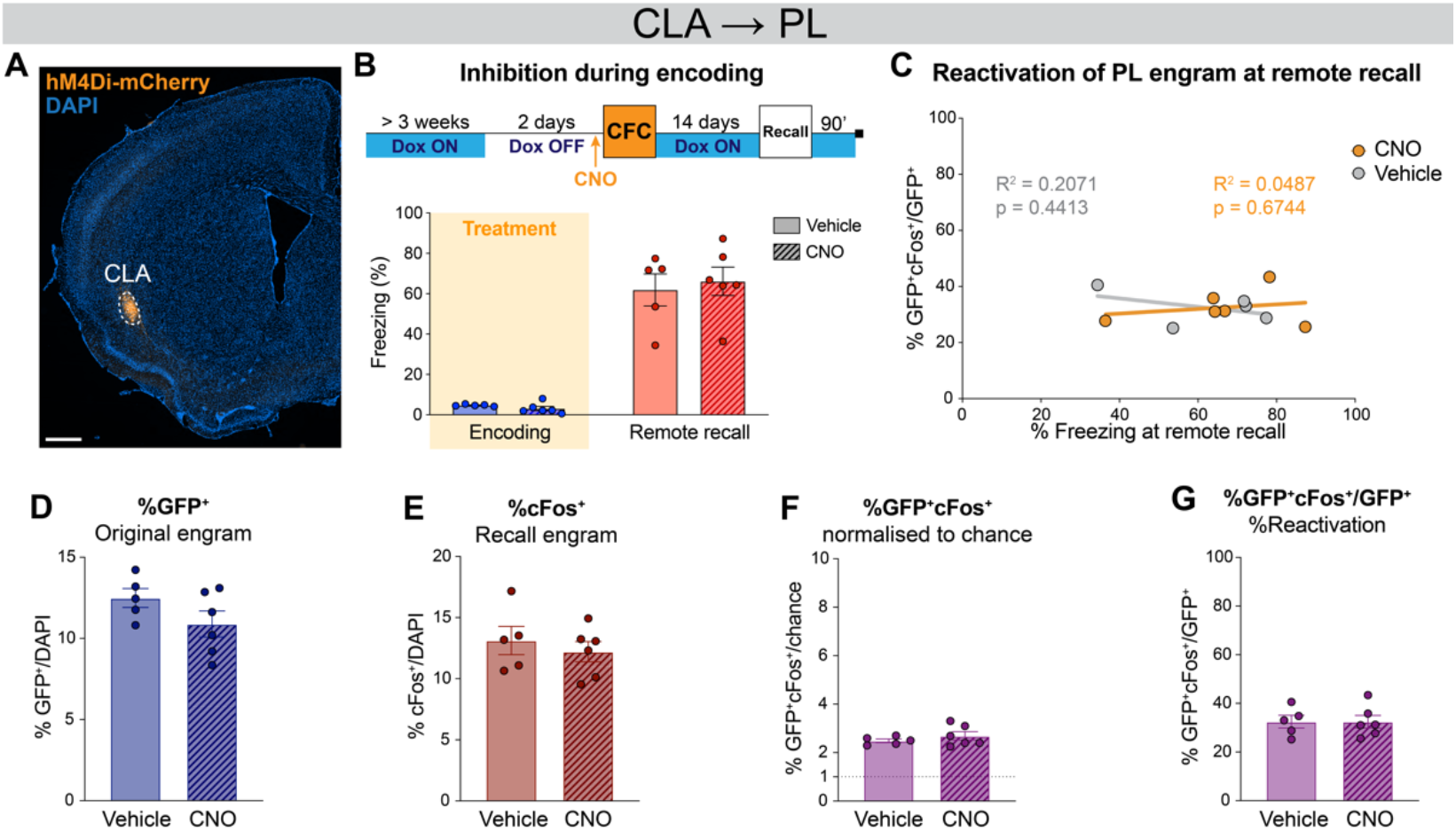
CLA → PL inhibition during encoding does not affect remote recall and engram reactivation in PL. **(A)** Representative image of the CLA input region. **(B)** Timeline and freezing percentage of CLA → PL inhibition during encoding and tested at remote recall. **(C)** Reactivation of PL engram cells (%GFP^+^cFos^+^/GFP^+^) at remote recall, correlated with freezing percentage for CNO (orange) and vehicle (grey) groups. Percentage of **(D)** GFP^+^, **(E)** cFos^+^, **(F)** double positives GFP^+^cFos^+^normalized to chance and **(G)** reactivation as %GFP^+^cFos^+^/GFP^+^. Correlation assessed with linear regression, R^2^ and p-value are reported on the graphs. n =5-6.

**Figure 2 - Table 1.**
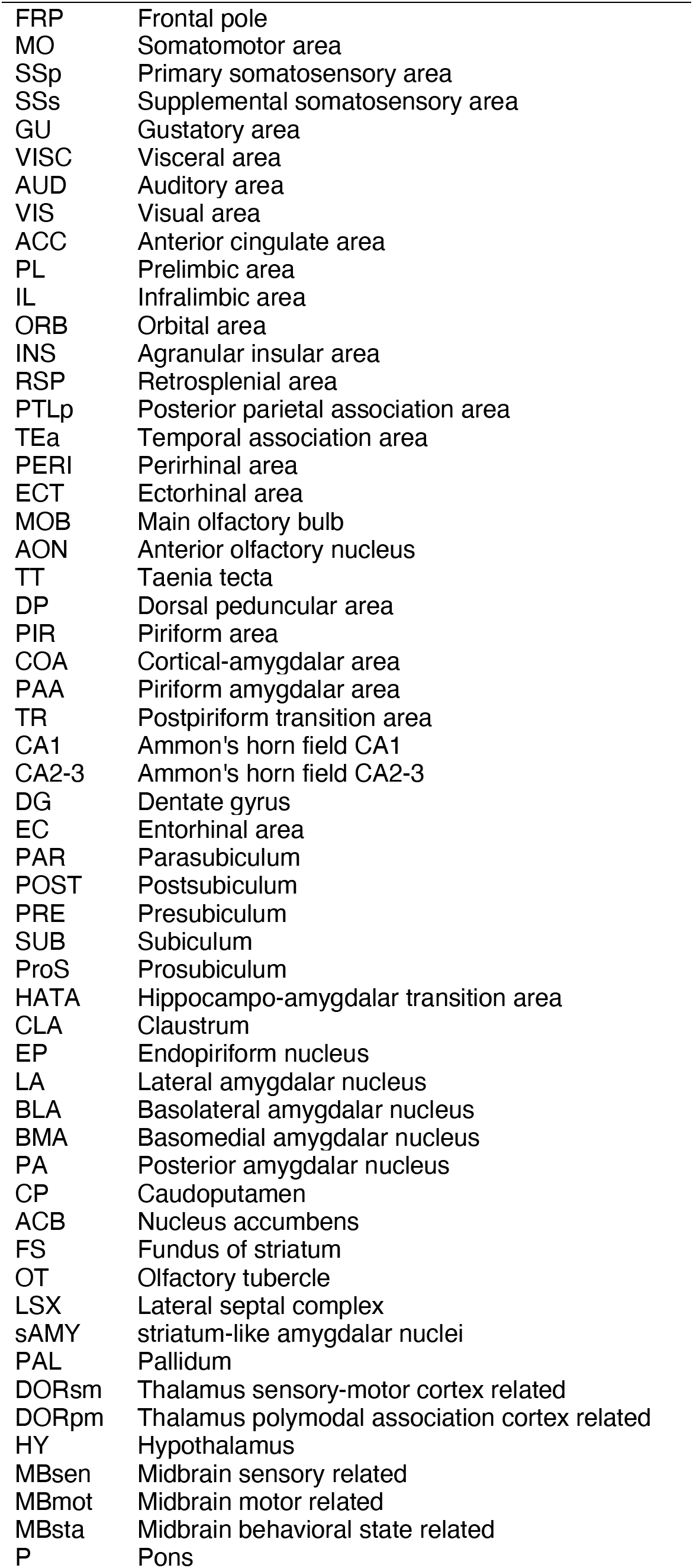
Abbreviations used for the brain regions

